# Population Genomics Informs Conservation Strategies for Critically Endangered *Kokia* Species in Hawai‘i

**DOI:** 10.1101/2025.07.17.665168

**Authors:** Weixuan Ning, Ehsan Kayal, Jonathan F. Wendel, Chuan-Yu Hsu, Zenaida V Magbanua, Olga Pechanova, Mitsuko Yorkston, Clifford W. Morden, Matthew J. Keir, Joshua R. VanDeMark, Daniel G. Peterson, Mark A. Arick, Corrinne E. Grover

## Abstract

Island endemic species are particularly vulnerable to extinction due to their limited geographic ranges and small population sizes. *Kokia* is a genus found exclusively in the Hawaiian Islands, whose species were once major components of local forests but have experienced significant population reductions due to habitat destruction and the consequences of invasive species. Although conservation of *Kokia* species has been an ongoing topic for over a century, records regarding historical efforts are sparse. Recently generated genomes for each of the three extant species provide the foundation for understanding genetic diversity and population structure for future conservation work. Whole genome resequencing of *K. cookei* (n = 23 samples), *K. drynarioides* (n = 92), and *K. kauaiensis* (n = 45) suggests that *K. drynarioides* has the lowest overall diversity, reflecting propagation from a limited part of the remaining gene pool, whereas *K. kauaiensis* exhibits the most diversity. Diversity in the primarily graft-propagated *K. cookei* is higher than expected, and slightly higher than in the free-living *K. drynarioides*. Notably, our analyses identified a source of novel variation in *K. cookei* in a cultivated plant historically labeled *K. drynarioides*. Population structure analyses reveal a single population for *K. cookei*, but three groups for each of the other two species. Importantly, our analyses identify clusters of related individuals, reflected in genetic distance and clustering metrics, which provide valuable information for increasing diversity in managed populations and in *ex situ* conservation collections. These results provide a genomic framework for ongoing efforts in restoring and maintaining diversity in these critically endangered Hawaiian species.

## 1. Introduction

The Hawaiian archipelago is the most remote major island chain in the world, renowned as a biodiversity hotspot with high levels of endemism due to adaptive radiations and geographic isolation (Barton et al., 2021; Cowie & Holland, 2008; Friedlander et al., 2020; Wall et al., 2015). Among the 1,381 native plant species, around 90% (1,238) are considered endemic (Rønsted et al., 2022), but many species now face extinction due to human activities. While human colonization began with the early seafaring Polynesians, extensive habitat modification happened as European colonists converted forests to farmland and introduced domestic animals and other invasive species (Boyer, 2008; Pejchar et al., 2020; Sakai et al., 2001). Over one third of native Hawaiian plant species are federally listed as endangered (Lim & Marshall, 2017; Rønsted et al., 2022; Werden et al., 2020) and over half are identified as “Species of Conservation Importance” by the Hawaiian Strategy for Plant Conservation due to their high extinction vulnerability combined with their cultural and/or habitat restoration value (*Laukahi Network - Hawaiian Rare Plant Species Coordination*, 2021; Weisenberger & Keir, 2014). Among these, 134 endemic plants are now considered extinct or extinct in the wild and nearly a quarter of native plant species are represented by fewer than 50 wild individuals (Werden et al., 2020; Wood et al., 2019).

The Hawai’i Plant Extinction Prevention Program (PEPP) is a conservation initiative dedicated to conserving Hawaiʻi’s most rare and endangered species, which currently lists 273 priority plant taxa that are either extinct, extirpated, or have fewer than 50 wild individuals remaining (*PEPPHI.org*, n.d.). Among these are all species within the genus *Kokia* (Malvaceae), with four described long-lived perennial flowering tree species endemic to the Hawaiian Archipelago (Figure 1). *Kokia* trees were once a common component of the Hawaiian xeric-mesic (i.e., dry to moderately moist) forests (Lewton, 1912; Sherwood & Morden, 2014); however, *Kokia* populations have plummeted since the early 1900s, mainly due to deforestation, competition from invasive plant species, and increased predation by grazing animals such as goats and cattle (Chynoweth et al., 2010; Hibit & Daehler, 2019). This decline has led to the extinction of one species (*K. lanceolata*) and the extirpation of a second (*K. cookei*) from the wild (Morden & Yorkston, 2018; Stone, 1967).

**Figure 1:**
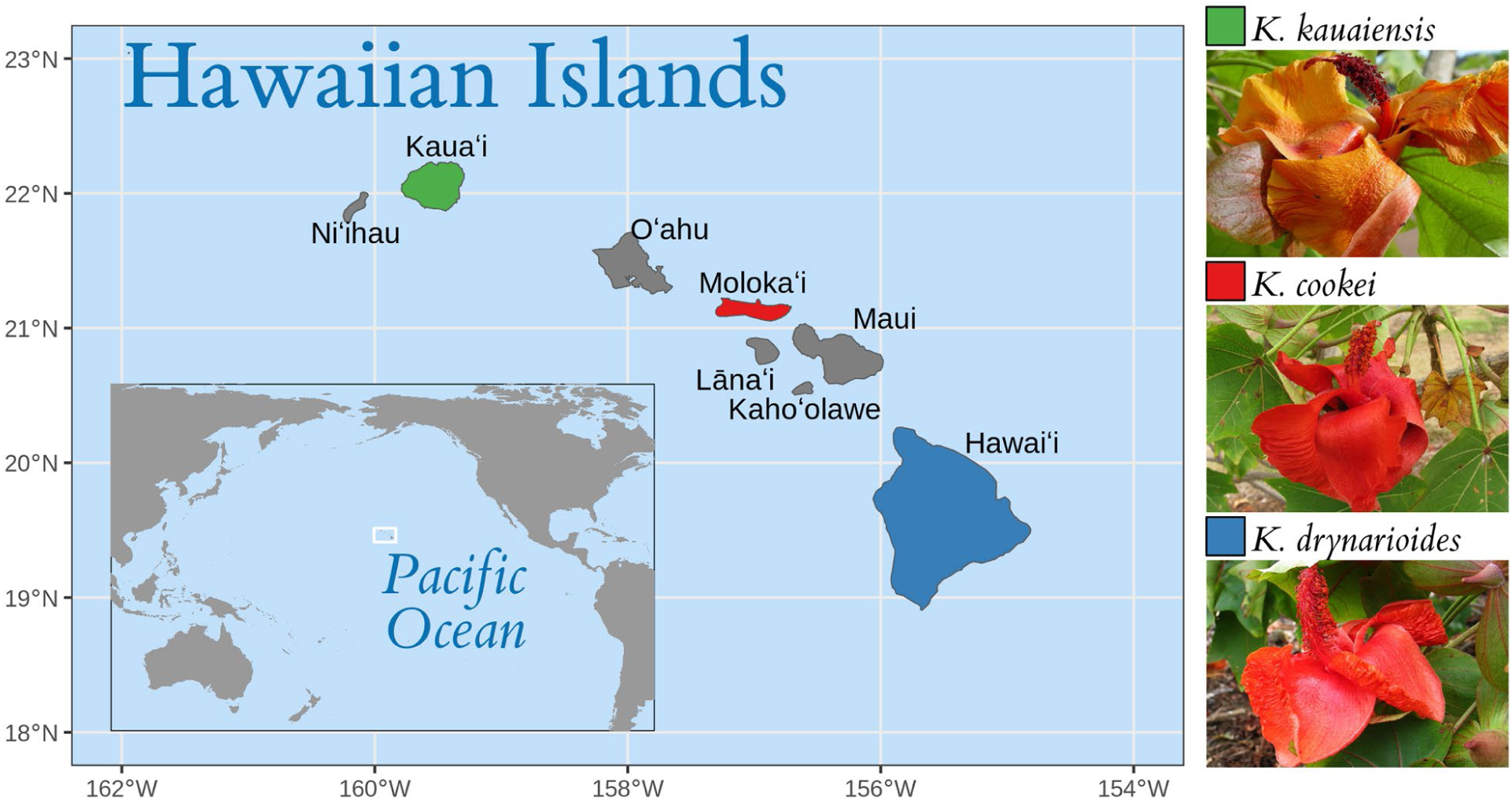
Native islands of the three extant species of Hawaiian highly endangered and endemic *Kokia* species (*K. cookei* Degener, *K. drynarioides* Lewton, *K. kauaiensis* Degener & Duvel). Map made with data from Natural Earth. Image credits: photos of *K. cookei* and *K. drynarioides* to David Eickhoff from Pearl City, Hawaii, USA (https://commons.wikimedia.org); photo of *K. kauaiensis* from the National Tropical Botanical Garden, https://ntbg.org/.

Adding to their vulnerability, each *Kokia* species is a single-island endemic whose fate has been tied to development on each island (Figure 1). Notably, the holotype of the extinct *K. lanceolata* Lewton was collected in the Wailupe Valley on O‘ahu (Hillebrand, 1995) only 5-10 years before its reported extinction (Rock, 1919). *Kokia cookei* Degener, endemic to Moloka‘i (Stone, 1967), narrowly escaped extinction by surviving in cultivation, first as free living descendants from a single wild tree (Degener, 1932a; Stone, 1967) but later only as grafts (on either *K. drynarioides* or *K. kauaiensis*) derived from a single cultivated individual (Sherwood & Morden, 2014) or from seeds derived from that individual or its surviving grafts.

The two remaining free-living species, *K. drynarioides* and *K. kauaiensis*, have fared slightly better, both still existing in the wild, albeit with varying success. Native to Hawaiʻi (colloquially, the Big Island), the remaining natural population of *K. drynarioide*s (Seem.) Lewton (native name “Hau hele ‘ula”) consists of fewer than 30 individuals, i.e., three mature trees and their offspring. Conservation efforts have been put in place to protect the remaining wild individuals, as well as increase the representation of *K. drynarioide*s on Hawaiʻi through intentional outplants (i.e., curating *K. drynarioides* seedlings to establish or increase managed populations). Currently, *K. drynarioides* has human-managed outplant populations in approximately 15 locations interspersed on Hawaiʻi Island; however, the genetic diversity represented by the remaining wild individuals, and thus the outplants, has been understudied.

Wild populations of *K. kauaiensis* (Rock) Degener & Duvel remain extant in nature on Kauaʻi, although they have also experienced recent declines. Population estimates for *K. kauaiensis* from circa 2000 estimated between 105 to 171 individuals, but the most recent survey (2022) suggests that only 19 wild individuals remain due to ongoing threats including landslides, fire, insects, and ungulates (U.S. Fish and Wildlife Service, 2022). Accordingly, managed populations of *K. kauaiensis* have been established in controlled areas with active threat management (e.g., fencing against grazing). Notably, this may not be the only decline experienced by *K. kauaiensis* populations. This species was considered nearly extinct by Degener (Degener, 1932b) in the early 20^th^ century and by subsequent botanists as “exceedingly rare” (Stone, 1967). The consequences of these population size fluctuations for diversity and population health in *K. kauaiensis* are unknown.

While conservation concerns for *Kokia* species became apparent shortly after their circumscription (Young, 1916), current understanding of intraspecific genetic diversity is limited. The only survey of genetic diversity in the genus was published a decade ago using 115 Random Amplified Polymorphic DNA (RAPD) markers, reported low intraspecific diversity, but noted it was within the range observed for other Hawaiian plant species (Sherwood & Morden, 2014). Unsurprisingly, given their current population sizes and distributions, *K. kauaiensis* exhibited the greatest number of polymorphic loci (52.3%), followed by *K. drynarioides* (32.2%) and finally *K. cookei* (16.5%). The authors suggested that, given their prevalence in remote, relatively undisturbed areas of Kauaʻi, diversity among *K. kauaiensis* individuals may be indicative of natural genetic variation in *Kokia* species in the absence of human activity or intervention (Sherwood & Morden, 2014). The remaining diversity in *K. drynarioides* and *K. cookei*, therefore, reflects their individual histories of population decline and genetic bottlenecks.

Although this initial research provided insight into diversity in *Kokia* species, it has been limited by the lack of application of modern genomic tools and resources. Recently, reference-quality genome sequences for all three extant *Kokia* species were published (Grover et al., 2017; Kayal et al., 2024), providing the foundation for improved genetic surveys. Leveraging these new genomes, we used recently collected and existing resources to collect genome-wide resequencing data from multiple specimens for each species: *K. cookei* (n = 23), *K. drynarioides* (n = 92), and *K. kauaiensis* (n = 45). Using these data, we evaluated genetic diversity and divergence within and among species. We identify groups of closely related individuals in each species and, for *K. drynarioides*, relate the composition of managed populations to these genetic groups. We hope that our results will inform conservation priorities by identifying genetically distinct individuals and populations suitable for propagation and management.

## 2. Material and methods

### 2.1 Sampling and sequence generation

In total, 163 *Kokia* samples were collected, including 57 existing DNA samples from the Hawaiian Plant DNA Library (HPDL; Morden et al. 1996) and 106 newly collected and silica-gel-preserved leaf samples. All samples were shipped to Mississippi State University (under permit I2665 to C.E.G.) for further processing. Initial sample identification based on historical records suggested that these samples represented 22 *K. cookei* (Kc), 93 *K. drynarioides* (Kd), and 45 *K. kauaiensis* (Kk) individuals (Table 1) with one *K. drynarioides* tree represented by four replicates (for a total of 96 *K. drynarioides* samples).

**Table 1:** *Kokia* specimens used in this study, partitioned by species, and range in sequencing coverage. General locations are given for wild trees, where available. Due to the protected nature of these species, some locations have been approximated or omitted. Unless otherwise specified, *K. drynarioides* samples are from Hawaiʻi and *K. kauaiensis* samples are from Kauaʻi. All *K. cookei* and *K. drynarioides* samples were provided by the HDPL (as noted) or by Hawai‘i State Department of Forestry and Wildlife (DOFAW) under permit I2665 to C. Grover, Iowa State University. Most *K. kauaiensis* samples were provided by the HDPL (Morden et al, 1996), and their collection locations are given. Samples noted with an * derive from the same tree, growing in the Hawaiʻi Volcanoes National Park (HAVO) greenhouse. These samples were from a tree historically considered *K. drynarioides*; however, our genomic analyses suggest this tree belongs to *K. cookei*.

| Species | Number of Samples | Location/Origin | Coverage (X) |
| --- | --- | --- | --- |
| *K. cookei* | 11 | D.T. Fleming Arboretum at Pu‘u Mahoe | 29.7 (27.2 - 33.9) |
|  | 8 | HDPL | 26.0 (20.2 - 36.7) |
|  | 2 | Olinda Rare Plant Facility | 29.0 (28.1 - 29.8) |
|  | 1 | private collection | 30.0 |
|  | 4* | HAVO | 66.0 (30.0 - 102.0) |
| *K. drynarioides* | 5 | Forest Bird Sanctuary Cabin Enclosure | 32.0 (21.0 - 43.0) |
|  | 20 | Hau‘aina | 24.0 (11.0 - 48.0) |
|  | 6 | Honomalino | 32.7 (29.0 - 42.0) |
|  | 2 | Ka'ūpūlehu makai | 30.8 (25.0 - 37.0) |
|  | 4 | Ka'ūpūlehu mauka | 38.5 (29.0 - 48.0) |
|  | 9 | Kīpuka Oweowe | 29.3 (23.0 - 42.0) |
|  | 8 | Koko Crater Botanical Garden | 32.9 (22.7 - 51.1) |
|  | 12 | Maka'ula 'o'oma | 26.2 (21.0 - 33.0) |
|  | 4 | Pu'u wa'awa'a cone | 26.2 (23.0 - 30.0) |
|  | 5 | Pu'u wa'awa'a exclosure | 29.2 (21.0 - 41.0) |
|  | 3 | remnant population | 32.1 (31.2 - 33.3) |
|  | 7 | Uhiuhi | 26.7 (18.0 - 38.0) |
|  | 6 | HDPL | 21.2 (16.9 - 27.0) |
|  | 1 | Iowa State University greenhouse | 37 |
| *K.kauaiensis* | 1 | Kalalau Valley population | 140.2 |
|  | 2 | Kawai'iki Canyon | 23.4 (22.6 - 24.1) |
|  | 2 | Koai'e Canyon | 21.6 (21.1 - 22.2) |
|  | 10 | Ku'ia Valley | 21.6 (16.3 - 25.2) |
|  | 24 | Pa'aiki Valley | 21.0 (10.2 - 31.8) |
|  | 1 | Pōhakuao | 15.6 |
|  | 5 | Waimea Botanical Garden, Waimea Falls | 23.4 (16.2 - 29.5) |

For DNA extraction, silica dried leaf tissue (30–50 mg per sample) was finely powdered using 2.0-mm ZR Bashing Beads (Zymo Research, Irvine, CA, USA) in a Geno/Grinder 2010 system (SPEX SamplePrep, Metuchen, NJ, USA) at 1,500 rpm for 30 seconds. Genomic DNA was extracted from the powdered leaf tissue using the Qiagen Plant DNeasy Mini Kit (Qiagen, Germantown, MD, USA) following the manufacturer’s protocol. DNA concentration and purity were assessed using a NanoDrop One spectrophotometer (Thermo Fisher Scientific, Waltham, MA, USA), and genomic DNA integrity was confirmed via agarose gel electrophoresis.

For Illumina DNA-Seq library preparation, 300–500 ng of genomic DNA per *Kokia* sample was processed using the NEBNext Ultra II FS DNA Library Prep Kit and NEBNext Multiplex Dual Index Primer Set (New England Biolabs, Ipswich, MA, USA) following the manufacturer’s instructions. AMPure XP beads (Beckman Coulter, Brea, CA, USA) were used to select the library size range (i.e. insert size range from 250 - 400 bp) using the double-sided size selection procedure. Additionally, 3-cycles of PCR were applied to the step that includes PCR-barcoding and enrichment of adaptor-ligated DNA. Each library was validated by using Agilent Bioanalyzer 2100 with Agilent DNA 1000 kit (Agilent Technologies, Santa Clara, CA) and quantified by using the Qubit fluorometer with the Qubit dsDNA HS assay kit (Life Technologies, Grand Island, NY). The prepared libraries were pooled in equimolar ratios and sequenced by Novogene Corporation (https://www.novogene.com/us-en/) as 150 bp paired-end reads using three S4 flow cell lanes on the NovaSeq 6000 system (Illumina, San Diego, CA, USA).

### 2.2 Interspecific genomic SNP calling

Given the similarity in morphological characteristics of three *Kokia* species and to first confirm all sampled individuals were correctly assigned to the right species using genetic data, we analyzed genetic relationships among the three *Kokia* species by mapping all sequenced 163 samples to the same reference genome: *K. kauaiensis* (Genbank ID: GCA_039519135.1) (Kayal et al., 2024). Raw reads from each sample were quality-filtered using Trimmomatic v.0.39 (Bolger et al., 2014)(ILLUMINACLIP:Adapters.fa:2:30:10:2:True LEADING:3 TRAILING:3 MINLEN:75) and then individually mapped to the reference genome using BWA v.0.7.17 (Li, 2013) with default parameters.

Genomic variation among the sequenced samples was identified using the DNA-Seq variant-calling pipeline from Sentieon Genomics v.202503-qgdzrem (Kendig et al., 2019). In brief, the BWA-mapped SAM file was converted into BAM format (‘util sort --sam2bam’). The PCR-duplicated reads within BAM file were identified (‘--algo LocusCollector’) and removed (‘--algo Dedup’), and surviving reads were realigned (‘--algo Realigner’) prior to haplotype calling (‘-- algo Haplotyper’). Finally, samples were jointly-genotyped (‘--algo GVCFtyper -- emit_all_sites’) by each individual species, emitting all sites, to produce three multi-sample VCF files that contain both variant and invariant sites. Each VCF was filtered via vcftools v.0.1.16 (Danecek et al., 2011) to keep: 1) only biallelic sites with no missing data (‘--max-missing-count 0 --max-alleles 2 --mac 2’) and 2) invariants sites (‘--max-maf 0’), both without insertions/deletions (‘--remove-indels’); all surviving sites were filtered by the average sequencing depth to remove the variation in low-coverage and high-copy genomic regions (‘-- min-meanDP 10 --max-meanDP 100’).

The VCF files for each of the three species (mapped to the same reference genome) were merged to generate a combined VCF and refiltered. We extracted the biallelic SNPs without missing data and with minor allele frequency > 0.001 (‘-m2 -M2 -i ’F_MISSING=0’ -q 0.001:minor’) in bcftools v.1.19 (Danecek et al., 2021; Li, 2011). Because diversity in genes is known to be important, we extracted genic SNPs using the annotation of the reference genome (Kayal et al., 2024) via vcftools (‘--bed’). All code, scripts, and parameters are available from https://github.com/Wendellab/KokiaResequencing.

### 2.3 Interspecific genetic variation comparison

Using the selected genic SNPs, we first calculated the proportions of observed heterozygous sites in each sample using vcftools (‘--het’) and the inbreeding coefficient (*F_IS_*) for each species. We performed statistical comparisons of heterozygosity and inbreeding levels between the three species using ‘pairwise.wilcox.test’ in R v.4.4.3 (R. Team, 2014). Then, fixed heterozygous sites across all 163 samples were removed using bcftools (‘--exclude “F_PASS(GT=’het’)=1”’) and the VCF was thinned using vcftools (‘--thin 100’) to include only one variant per 100 bp window. Genetic relationships between all 163 sequenced samples were estimated in Plink v.1.9 (Purcell et al., 2007), using Principal Component Analysis (PCA), and the first 20 PCs were calculated (‘--pca 20’). Using the same set of independent SNPs, we evaluated the population structure of 163 samples using the R package LEA (Frichot & François, 2015). Specifically, the number of population groups (*K*) was estimated from 1 to 15, with each *K* replicated 10 times. The best model (i.e., *K*) was selected based on the lowest cross-entropy value.

Because the four samples from the Hawai’i Volcanoes National Park (HAVO) are technical replicates (Table S1), only one (HAVO_1A) was used. Additionally, due to its reassignment to *K. cookei* (see Results), HAVO_1A was analyzed with the remaining *K. cookei* (see Results) rather than its original species assignment (*i.e., K. drynarioides*). Genic sequence diversity and genetic differentiation between the three species were compared via Pixy v.1.2.10.beta2 (Korunes & Samuk, 2021). Using the combined VCF containing both variant and invariant sites from all three species, we first extracted genic region sequences via vcftools (‘--bed’) with the *K. kauaiensis* reference genome. The average nucleotide sequence diversity (*pi*/𝝅𝝅) within genic regions was calculated for each species, using a sliding-window size of 10 kb. Similarly, pairwise sequence divergence (*dxy*) between three species was also calculated in Pixy using the same window size. Again, HAVO was represented by only one replicate (*i.e.*, HAVO_1A). We partitioned the results for each chromosome and plotted the figure using ggplot2 (Wickham & Sievert, 2009).

### 2.4 Intraspecific genomic SNP variant calling and analysis

To provide a better resolution for intraspecific variation, we mapped the sequencing reads from each sample to its own species reference genome. Specifically, similar to the steps and setting as above, the trimmed reads of 22 *K. cookei* plus one HAVO sample (HAVO_1A) were individually mapped to the *K. cookei* reference genome (GeneBank ID: GCA_039519145.1), all remaining 92 *K. drynarioides* were mapped to *K. drynarioides* reference genome (GeneBank ID: GCA_002814295.2), and all 45 *K. kauaiensis* were mapped to the *K. kauaiensis* reference genome. We used Sentieon to genotype the variants and invariant genetic sites within each sample (i.e., gVCF), and jointly-called the variants across all samples within each species, respectively, to produce three multi-sample VCF files. The variant and invariant sites in each VCF file were filtered using the same threshold as above.

For the three intraspecific variant VCF files, sites where heterozygosity was fixed (i.e., all samples are heterozygous) were removed as uninformative, and the resulting files was filtered for only genic variants using the predicted annotation files for each species, as above. We first calculated three pairwise genetic distance matrices using the R packages vcfR v.1.15.0 and ape (‘dist.gene’ function) , to estimate the relatedness between samples in each species. To investigate the population genetic relationships, first overrepresented closely-linked SNPs were removed using vcftools (‘--thin 100’). These independent SNPs were used to calculate: 1) the first 20 PCs for each species in Plink; 2) the population genetic structures in each species via LEA, with *K* = 1 to 15, and each *K* was replicated 10 times. Moreover, we repeated the above analysis for *K. cookei* by removing the HAVO sample from the variant VCF file via bcftools (subset ‘-S’), keeping the same filtering criteria above. Finally, we quantified: 1) the total genetic diversity (𝝅𝝅), and 2) the amount of genetic diversity within each sub-groups/populations, for each species via pixy, with the same sliding-window size as above.

## 3. Results

### 3.1 Sequencing depth and filtering

We generated an average of 58 million reads (ranging 23-273 M) per sample, corresponding to an average of 28X coverage resequencing (ranging 10X -140X), for the 163 *Kokia* samples representing all three extant species: *K. cookei* (Kc; 22 individuals), *K. drynarioides* (Kd; 93 individuals), and *K. kauaiensis* (Kk; 45 individuals; Table 1). To facilitate interspecific comparisons, we mapped all samples to a common reference. Preliminary analyses indicated that using either *K. drynarioides* or *K. kauaiensis* as a common reference for all species produced similar results with respect to read mapping, coverage, and PCA-based sample clustering (Table S1). Using *K. kauaiensis* as a common reference, we recovered ∼489.9 million (M) sites passing initial filtering for each genome (genome size: ∼520 Mbp (Seelanan et al., 1997)), representing >93% of the genome per sample (Table S2). Notably, only 0.45 - 1.32% of sites were variable within each species, indicating high homogeneity among samples. Because diversity in genes is important for conservation, we further focused our analyses on gene regions only, resulting in ∼86.6 M sites (Table S2). Variation within genes was even lower, ranging from 0.15 - 0.55% of total genic sites.

### 3.2 Species separation and identification

We first evaluated the divergence between species to provide a foundation for comparing diversity among species. Principal component analysis (PCA) of genome-wide genic SNPs segregated the three extant *Kokia* species (Figure 2A), with the two highest eigenvectors explaining over 50% of the variation (PC1 = 33.78% and PC2 = 16.45%). *Kokia drynarioides* and *K. cookei* appeared more similar to each other than either was to *K. kauaiensis*, primarily separating along the second eigenvector (PC2), whereas *K. kauaiensis* was distinct along PC1. Notably, all four samples taken from the single tree in cultivation at the Hawaiʻi Volcanoes National Park (HAVO), which was historically considered *K. drynarioides*, were grouped with the *K. cookei* samples rather than with *K. drynarioides*. LEA analysis further confirms the HAVO tree as *K. cookei*, as these samples were indistinguishable from other *K. cookei* samples (Figure 2B). Overall, the LEA analysis suggests seven groups within these three species: a single, nearly uniform group for *K. cookei*, which includes HAVO, and three groups each for *K. drynarioides* and *K. kauaiensis* (Figure 2B). Among the *K. drynarioides* and *K. kauaiensis* populations, genetic structure ranges from nearly uniform samples (e.g., the 65 *K. drynarioides* samples composed largely of P5; Figure 2B) to samples with highly mixed ancestry (e.g., Kd_3149, and Kk_19S8; Figure S1).

**Figure 2:**
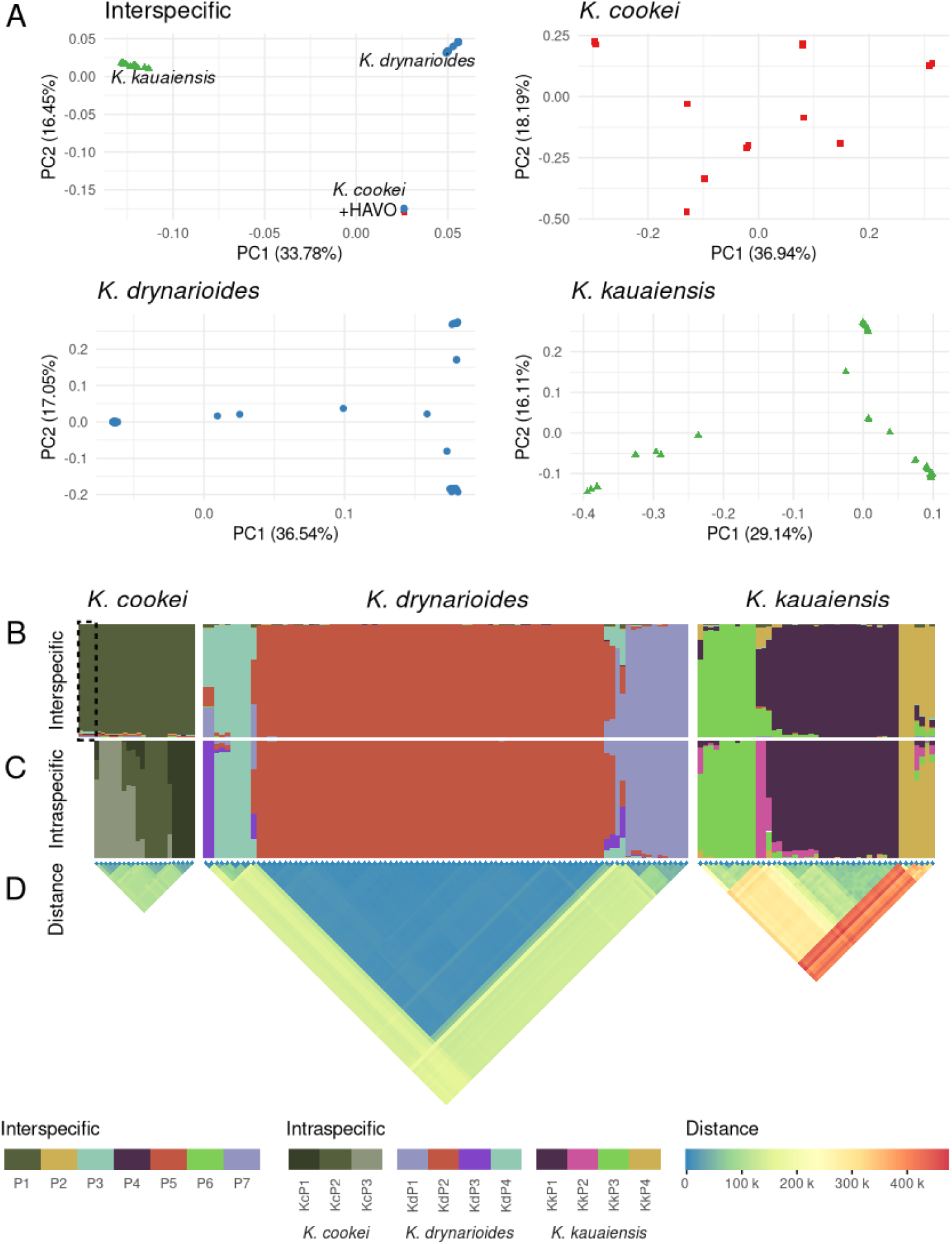
Relationships among species and individuals in three *Kokia* species. (A) PCA among all samples (mapped to the *K. kauaiensis* genome) and within species (mapped to species-specific genomes) using variant genic sites and including all four HAVO replicates. Each species is distinguished by color and shape: *K. cookei* (red square), *K. drynarioides* (blue circle), and *K. kauaiensis* (green triangle). In the interspecific panel, blue circles grouping with *K. cookei* represent the four HAVO replicates, as noted in the PCA. Percent of variance explained by each axis is noted. (B-C) Ancestral population structure (or genetic ancestry) for interspecific (*K_Int_*=7) and intraspecific (*K_KC_*=3; *K_Kd_*=4; *K_Kk_*=4) analyses. Colors are retained across B and C to the extent possible. HAVO samples are shown only in the interspecific analysis and are noted by a dashed box. (D) Genetic distance (in number of variant sites) among individuals based on the intraspecific analyses.

### 3.3 Interspecific genomic comparisons highlight genetic variation in endangered species

Because the HAVO samples grouped with *K. cookei* in both the PCA and LEA analysis, we removed these samples from the *K. drynarioides* sample set and included one representative (HAVO_1A) in the *K. cookei* sample set for the remaining analyses. This reduced the number of genic variant sites within *K. drynarioides* from 276,423 to 169,309 (Table S2) and increased the number of genic variant sites within *K. cookei* from 71,574 to 105,563. Using this set of SNPs, we calculated diversity within each species by chromosome. As expected, genetic diversity (Figure 3A) in all species is low, being orders of magnitude lower than outcrossing species (e.g., maize (Dominguez et al., 2024)), and an order of magnitude lower than found in the related cotton genus (Ning et al., 2024; Yuan et al., 2021). Genetic diversity in *K. drynarioides* (*π*=5.32 x 10^-4^) is slightly lower than in *K. cookei* (*π*=6.28 x 10^-4^), perhaps due to the large number of very similar individuals sampled or perhaps due to the similarly severe, recent population reduction experienced by *K. drynarioides*. As expected from historical census numbers, *K. kauaiensis* has the highest genetic diversity (*π*=1.29 x 10^-3^), perhaps indicating something closer to the level of diversity in *Kokia* species that might have experienced less drastic population reductions. Notably, this comparative analysis suggests that, although *K. cookei* diversity is low, it is not radically different from its congeners.

**Figure 3:**
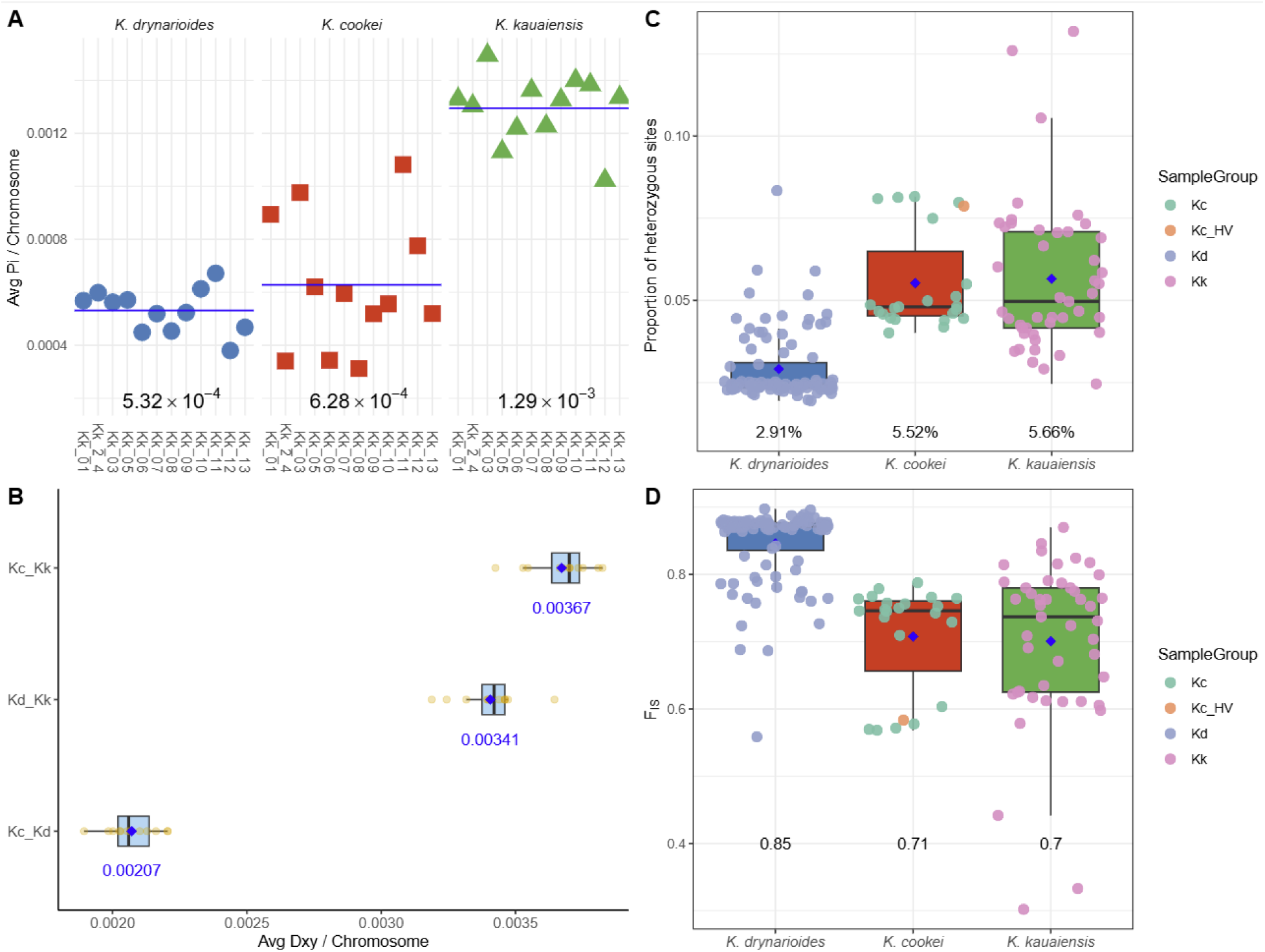
Genetic diversity and divergence among *Kokia* species. (A) Average nucleotide diversity (*π*) per chromosome for *K. drynarioides* (blue), *K. cookei* (red), and *K. kauaiensis* (green); horizontal lines indicate the mean per species. (B) Average pairwise divergence (*dxy*) per chromosome among species pairs; boxplots show distribution with blue diamonds indicating the mean, which is also listed below each boxplot, and black lines to indicate the median. (C) Proportion of heterozygous sites (*He*) per individual across species, with means labeled below each group. (D) Inbreeding coefficient (*F_IS_*) per individual, grouped by species. In all analyses, one HAVO was included as *K. cookei*. In both (C) and (D), the HAVO individual (HAVO_1A; Kc_HV) identified as *K. cookei* is indicated as an orange circle, and the remaining species are colored as indicated.

Given this low diversity, it is unsurprising that heterozygosity (*He*) within each species was also low (2.91 to 5.66%) and inbreeding (*F_IS_*) values were relatively high (0.70 to 0.85; Figure 3). Both *He* and *F_IS_* were variable among accessions in each species, albeit with a tendency to cluster more in *K. cookei* and *K. drynarioides*. In *K. kauaiensis*, the only remaining natural population, estimates of *He* and *F_IS_* were distributed around the mean, with a few outliers that likely represent the gene flow captured by the LEA analysis. Both *K. cookei* and *K. drynarioides*, however, were somewhat different, having a small proportion of individuals exhibiting higher *He* and lower *F_IS_* while the remaining individuals exhibited elevated *F_IS_* and low *He.* Interestingly, although *π* was similar between *K. cookei* and *K. drynarioides*, *He* was statistically lower and *F_IS_* was statistically higher in *K. drynarioides* relative to the other two (pairwise Wilcoxon rank-sum *P* <0.01; Figure 3). This likely reflects several phenomena, including the large number of highly similar samples in *K. drynarioides*, along with the fixed heterozygosity arising from those *K. cookei* samples that are clones of a single original tree.

### 3.4 Fine-scale population structure analysis of each species to guide future conservation efforts

Because all three species of *Kokia* are endangered, we also analyzed each species separately to better inform individual conservation efforts. Variants were called for each species using its own respective genome and annotation, which, as expected, yielded fewer variant sites (no missing data; Table 2). This was particularly apparent in *K. drynarioides*, which had an ∼50% reduction in the number of variant SNP sites.

**Table 2:** Number of individuals and sites evaluated in the intraspecific analyses. Because the HAVO tree appears to represent novel diversity in *K. cookei*, values are listed when both including or excluding the HAVO sample. Nucleotide diversity (𝝅𝝅) is listed for the species overall, and for each LEA-identified subpopulation (Figure 2). For *K. kauaiensis*, these correspond to natural populations consistent with geographic locations; for the two managed species, individuals were assigned based on their majority population structure as defined by LEA analyses.

| Species<br>(Same sp.<br>reference) | Sample<br>size | Total sites<br>surveyed | Variant<br>sites | Variant<br>sites<br>(genes) | $\pi$<br>overall | $\pi$<br>Pop1 | $\pi$<br>Pop 2 | $\pi$<br>Pop3 | $\pi$<br>Pop 4 |
| --- | --- | --- | --- | --- | --- | --- | --- | --- | --- |
| *K. cookei* +HAVO | 23_<br>22† | 82.9 M | 1.3 M | 128,355_<br>112,463† | 6.89e-4_<br>6.73e-4† | 5.62e-4† | 7.28e-4† | 5.93e-4† | - |
| *K. drynarioides* | 92_ | 86.6 M | 2.6 M | 172,777_ | 4.00e-4_ | 2.73e-4_ | 3.87e-4_ | 1.45e-3_ | 4.43e-4 |
| *K. kauaiensis* | 45_ | 85.2 M | 6.2 M | 465,146_ | 1.29e-3_ | 1.48e-4_ | 5.86e-4_ | 5.48e-4_ | 7.31e-4 |

† excluding HAVO

PCA analyses within each species revealed both tight clusters of highly similar individuals interspersed with loose clusters and more dissimilar individuals (Figure 2A), although this was less apparent in *K. cookei*, particularly when HAVO was excluded (Figure2, Figure S2). The variance described by the first two principal components was very similar for *K. cookei* (36.94% and 18.19%) and *K. drynarioides* (36.54% and 17.05%), and lowest in the more diverse *K. kauaiensis* (29.14% and 16.11%; Figure 2A). Population structure analysis of each species suggests that each species may contain 3 or 4 populations or genetic groups, as suggested from the PCA results. Details for each species, which we hope may inform conservation efforts, follow.

#### 3.4.1 K. cookei

PCA for the *K. cookei* samples loosely grouped samples into three main clusters of five samples, a smaller cluster containing two samples, and an additional five samples were interspersed (Figure 2). LEA analysis yielded results largely congruent with the PCA, identifying the three larger clusters (i.e., upper left, middle, and upper right) as homogeneous samples (KcP1-KcP3; Figure 2), and the interspersed samples as mixtures of these three populations. Although records of provenance are sparse, inferences regarding historical propagation can be gleaned from these data. Samples Kc_A_2/5/7/8 and Kc_3156 (Figure S1) appear to all derive from one graft lineage, whereas Kc_A_3/6/9/11 and Kc_3157 (Figure S1) may all derive from a different graft lineage. In both cases, the trees are thought to derive from seedlings originating from an early *K. cookei* graft (or airlayer clones of those seedlings). The third homogeneous group contains Kc_K_20 which is a graft-clone kept in a private collection on Maui, as well as Kc_A_1_1, which may be derived from a seedling of the original graft, and HPDL samples Kc_3148/3159/3160. The remaining samples (Figure 2) exhibit signatures of those three primary lineages and may derive from offspring generated by one or more graft lineages. Among these are the HDPL samples (i.e., Kc_3148, Kc_3156-3160, and Kc_4372-4373), Kc_A_4 (graft-derived offspring), Kc_A_10 (unknown provenance), and Kc_F1 (in cultivation on O‘ahu).

As expected, distance among individuals within each homogeneous group is relatively low. Genetic differences ranged from an average of 40.7 - 46.2 thousand (k) alleles (∼18% of the total) within groups to an average of 103.0 - 105.2 k differences (∼46%) between groups. Diversity estimates among these *K. cookei* samples (𝝅𝝅 = 6.73 x 10^-4^) was still exceptionally low and marginally increased when the reassigned HAVO sample was included (𝝅𝝅=6.89 x 10^-4^; Table 2). PCA of the *K. cookei* data + HAVO (Figure S2) shows a clear distinction between the HAVO sample and the remaining *K. cookei* samples, separating these samples on PC1 (explaining 41.1% of the variance) and exhibiting separation among the graft-derived samples only on PC2 (22.1% of variance). Together, the stark reduction in diversity when excluding HAVO and the clear distinction of HAVO on the intraspecific PCA suggest that this plant may be a novel source of diversity in this depauperate species

#### 3.4.2 K. drynarioides

PCA for the *K. drynarioides* samples (PC1 = 36.54%; PC2 = 17.05%) exhibited three apparent clusters and several interspersed samples; however, LEA identified an additional, fourth group not apparent in the interspecific LEA or the PCA (KdP1-KdP4; Figure 2B, 2C). Unlike *K. cookei*, where a majority (68%) of samples had an admixture coefficient > 0.999 (i.e., were assigned to a single ancestral group), only 45 of the 92 *K. drynarioides* samples (49%) exhibited a homogeneous genetic signature (Figure 2C). Most of these homogeneous samples (34) were assigned to KdP2, a genetic group containing 75% of the *K. drynarioides* samples (i.e., 69 of 92) and corresponds to the large cluster of samples near the left axis on the PCA (Figure 2A). The three *K. drynarioides* individuals originating from the remnant population (i.e., the last wild population on Hawaiʻi) are placed within the same population (KdP1), which is notably the only natural population in the survey of this species. Because most *K. drynarioides* trees are outplants, and are therefore managed under human direction, group membership for the remaining trees instead reflects a history of shared genetic ancestry. When plotted by location (Figure S3), it is apparent that KdP2 has been successfully incorporated into all outplant locations, whereas the remaining groups (KdP1, KdP3, and KdP4) have limited spread amongst outplant populations.

Genetic distances among *K. drynarioides* individuals is larger than that of *K. cookei* when comparing individuals between groups; however, KdP2 is notably uniform, exhibiting an average of 9,404 alleles that distinguish individuals compared to the overall average of 69,486 alleles. LEA groups are also generally recapitulated by the absolute genetic distance, although some substructuring of KdP1 may be apparent. Diversity estimates for *K. drynarioides* remained low (𝝅𝝅=4.00 x 10^-4^) and were somewhat lower than the interspecific estimate (𝝅𝝅=5.32 x 10^-4^), which notably included the subsequently reassigned HAVO sample. Within LEA groups, diversity varied from 𝝅𝝅=1.50 x 10^-5^ in KdP3 to 𝝅𝝅=4.43 x 10^-4^ in KdP4 (Table 2). Because only two samples were placed in KdP3, it remains uncertain whether those samples reflect a pocket of novel diversity or if the number of reconstructed populations is an overestimate.

#### 3.4.3 K. kauaiensis

Similar to *K. drynarioides*, PCA for *K. kauaiensis* revealed three apparent clusters and some interspersed individuals (Figure 2A) that are largely consistent with their collection location (Table S1). As with *K. drynarioides*, the intraspecific LEA (Figure 2B, 2C) recovered an additional, albeit small (2 samples) group, although the remaining groups were more even in size. Because most *K. kauaiensis* samples were taken from natural populations on Kauaʻi (circa 2000), each LEA population generally corresponds to a geographic location. KkP1 contains almost all of the Paʻaiki Valley samples except for a single sample (Kk_3177), which was collected from Paʻaiki Valley, but exhibits genetic signatures from KkP3, which is generally found in the Ku‘ia Valley. KkP4 is composed of trees located in Koai‘e Canyon, Kawai‘iki Canyon, and Pōhakuao; the first locations separated by less than 3 km and both of which are approximately 12 km from Pōhakuao. The smallest group, i.e., KkP2, is composed of a single outplant in the Kalalau Valley and a Waimea Falls (cultivated) accession (Table S1). The four remaining Waimea Falls samples exhibit genetic signatures suggestive of their origins: two (i.e., Kk_3152 and Kk_3154) likely originate from the Koai‘e Canyon, Kawai‘iki Canyon, and Pōhakuao population (KkP4), and the other two (i.e., Kk_3151 and Kk_3153) from the Pa‘aiki Valley population (KkP1). Considering Waimea Falls represents a cultivated collection, it is not surprising the samples are from multiple, seemingly unrelated, populations. While no Waimea sample was assigned to the Ku‘ia Valley population (KkP3), Kk_3151 exhibits a mixed genetic signature from all four populations, perhaps indicating that it is the offspring of other cultivated plants from different original sources.

In general, *K. kauaiensis* individuals exhibit less uniformity than the other two species, consistent with its recent history of larger population sizes (although see Discussion). Notably, only 18 samples (40%) displayed a single genetic signature, whereas the remainder exhibited signatures of more than one population (Figure 2C), suggesting either gene flow or some recent, shared ancestry. Accordingly, absolute genetic distance among individuals was higher in *K. kauaiensis* than the other two *Kokia* species (Figure 2D), averaging 127,752 alleles within population and 344,038 alleles among populations. Within population diversity in *K. kauaiensis* was varied approximately 2.5-fold from 𝝅𝝅=5.48 x 10^-4^ in KkP3 to 1.48 x 10^-3^ in KkP1 (average 𝝅𝝅=1.29 x 10^-3^).

## 4. Discussion

Similar to myriad other Hawaiian species, trees in the genus *Kokia* have become a casualty of the arrival of humans to the archipelago. Now highly endangered, *Kokia* species were once a common feature of the Hawaiian xeric-mesic forests (Sherwood & Morden, 2014), radiating amongst the Hawaiian Islands (Kayal et al., 2024; Sherwood & Morden, 2014) and resulting in four species, only three of which remain today (i.e. *K. cookei*, *K. drynarioides*, and *K. kauaiensis*). Because the history of human activities varied among the islands, the resulting population declines have led to differing outcomes for each species. While *K. kauaiensis* has maintained continuity of natural populations, *K. drynarioides* was reduced to a natural population consisting of only three adult trees and their offspring and *K. cookei* was saved from a single remaining tree to persist as a propagated graft (Wood et al., 2019) or as offspring from that original graft (although see discussion below).

Despite the importance of these species to the Hawaiian forests and their critically endangered status, little was known about the persistence of genetic diversity within and among species aside from a single survey using ten random amplified polymorphic DNA (RAPD) markers, which found a surprisingly high degree of polymorphism for all three species, albeit with limited sampling for *K. cookei* (6 individuals) and *K. drynarioides* (5 individuals) (Sherwood & Morden, 2014). Our results extend this previous research using whole genome resequencing (>10X coverage) for an expanded sample set: *K. cookei* (n = 22), *K. drynarioides* (n = 93, including one of uncertain status), and *K. kauaiensis* (n = 45). Importantly, our results leverage both interspecific and intraspecific analyses, the former allowing us to identify a misidentified tree and compare outcomes for each *Kokia* species, and the latter describing population dynamics in each species and laying the foundations for more informed conservation efforts.

### 4.1 Genetic diversity of Hawaiian endemic Kokia spp

Because the three extant *Kokia* species originate from different islands in the Hawaiian archipelago, each has been subjected to different external pressures (e.g., habitat loss, invasive species proliferation, etc.). Accordingly, these species may have experienced differences in population reductions and loss of diversity. We first gauged the relative relationships among species by mapping all samples to a common reference genome (i.e., *K. kauaiensis(Kayal et al., 2024)*, finding a high rate of interspecific mapping (>99% of reads; Table S1), which is unsurprising given their recent divergence time and low synonymous substitution values (Kayal et al., 2024). As expected from previous analyses (Sherwood & Morden, 2014), genetic diversity is low in each species, resulting in only 720,546 variant nucleotide positions out of 86.6 M genic sites. Among species, *K. kauaiensis* exhibited the most diversity, reflecting a historical census from circa 2000. Since that time, however, *K. kauaiensis* populations have experienced a dramatic reduction in population size, from about 145 - 170 trees in 1996 to only ∼19 trees in 2022 (U.S. Fish and Wildlife Service, 2022). Without additional sampling, the consequences of this significant population reduction for diversity are unknown; however, the current analyses provide a framework for understanding historical diversity and incorporating that information into the conservation goals, particularly in sourcing trees for establishing outplant populations.

Because both *K. drynarioides* and *K. cookei* experienced more drastic and prolonged bottlenecks than did *K. kauaiensis*, they naturally exhibited lower levels of genetic diversity (π=5.32 x 10^-4^ and π=6.28 x 10^-4^ versus 1.20 x 10^-3^, respectively); however, based on the previous RAPD analysis, it was unexpected for them to exhibit similar levels of genetic diversity. Interestingly, not only was diversity in *K. drynarioides* slightly lower than in *K. cookei* (Figure 3), but it also exhibited lower average heterozygosity, as well as a higher inbreeding coefficient (Figure 3).

These observations are challenging to explain but likely result from a combination of recent divergence between the two species (Figure 3), significant recent inbreeding in *K. drynarioides*, and fixed heterozygosity among grafts. Notably, although *K. cookei* populations have remained small since its extirpation, diversity within this species is similar to its congeners, suggesting that there exists enough genetic diversity to establish outplant populations in protected areas and raises hope for conservation efforts in species with very small starting populations.

Importantly, this comparative analysis also identified a tree from the Hawaiʻi Volcanoes National Park (HAVO) that has historically been considered *K. drynarioides*, but whose variants (>94%) were overwhelmingly associated with *K. cookei* (Figure 2). Originally, two samples from this single tree were sequenced (i.e., HAVO-1a/b), and the tree was later resampled (i.e., HAVO1A/B) to confirm the findings. While long presumed *K. drynarioides*, the origins of the tree are uncertain, although it may originate from seeds sent by George P. Cooke to Joseph Rock circa 1957 (Woolliams & Gerum, 1992). This tree was originally planted in the HAVO National Park more than half a century ago (Norris, 1967; Woolliams & Gerum, 1992), where it was later rescued from an emergent root sucker after the mature tree was destroyed by a falling ʻŌhiʻa (*Metrosideros polymorpha*). Local knowledge within DOFAW notes that multiple species of *Kokia* were planted in various places around Hawaiʻi at various times, although the origins and identities of many of these plants have been lost. The *K. cookei*-dominant genetic signature of the HAVO samples suggests that this tree should be reassigned to *K. cookei* and maintained as such hereafter.

### 4.2 Genetic diversity among K. cookei

*Kokia cookei* is the most critically endangered species in the genus, surviving entirely in cultivation as single-source grafted clones or seeds from those clones and making it effectively a population of one (Sherwood & Morden, 2014), aside from the newly identified tree from the HAVO National Park. Accordingly, nucleotide diversity (𝝅𝝅) in *K. cookei* is low, albeit comparable to its entirely free-living congener *K. drynarioides*, suggesting that population recovery in this species could be similar. Importantly, the identification of a novel source of *K. cookei* variants both provides additional diversity for this species but also suggests that other historically planted trees with uncertain provenance may obscure additional sources of novel diversity through misidentification. Furthermore, this additional genetic material will likely benefit conservation efforts in this species by including additional diversity in these managed populations (Figure S2).

### 4.3 Restoration of K. drynarioides populations

Conservation efforts in *K. drynarioides* have largely focused on increasing population size and locations via outplants on Hawaiʻi. Although the natural population of *K. drynarioides* have been reduced to only between two to six wild trees, conservation efforts have established over 1,300 plants in 7 locations around Hawaiʻi (US Fish and Wildlife Service, 2020). Such an extreme bottleneck, however, puts *K. drynarioides* at high risk of inbreeding depression, which can result in vulnerability to disease and reduced adaptability in the face of environmental changes. Since the 1950s, seeds have been collected in an effort to restore *K. drynarioides* populations in the wild, mainly through outplanting and natural restoration projects, some of which have yielded natural recruits (Libby et al., 2022).

Despite this concerted effort, conservation in *K. drynarioides* has been operating in the absence of genetic data to guide plantings. Our analyses provide a genetic context for managing diversity in *K. drynarioides*. The data suggest that most *K. drynarioides* samples are highly genetically homogeneous and exhibit very high levels of inbreeding (*F_IS_* = 85%; Figure 3), which is a natural consequence of prolonged, significant population size reductions and subsequent human-mediated propagation from the few remaining individuals. Unsurprisingly, all three wild specimens (Kd_3041, Kd_3042, and Kd_3043) belonged to the same ancestral population and are largely homogeneous in our LEA analysis (Figure 2B-C). More importantly, however, is the identification of distinct groups of *K. drynarioides* whose genetic distance and population reconstruction suggests more diverged ancestry and which are underrepresented in current outplant populations (Figure S3). These groups provide the basis for establishing diverse populations, allowing DOFAW and PEPP to include the least related individuals in subsequent outplant populations, thereby increasing genetic diversity in managed populations.

### 4.4 Reviving populations of K. kauaiensis

Of the three extant *Kokia* species, *K. kauaiensis* has historically had the most robust natural populations, typically surviving around slopes of the remote valleys of western Kauaʻi Island (Sherwood & Morden, 2014). Although these areas have remained largely human-inaccessible, competition from or predation by invasive species remains a problem. Escape and occupation by feral ungulates, namely goats, pigs, and deer, have had dramatic negative impacts on native plant populations (Leopold & Hess, 2017), as have other herbivores (e.g., nonnative rodents and insects). Because predation pressure by opportunistic herbivores is not unique to *Kokia*, concerted efforts at regulation and eradication of nonnative species has had a positive impact on recovery of some native Hawaiian plant populations, although reports of these invasive species persist (Chynoweth et al., 2010; Hess et al., 2017). In the most recent review of the status of *K. kauaiensis* (U.S. Fish and Wildlife Service, 2022), predation threats were listed as prominent, as were impacts by invasive plant species (competitors) and habitat destruction (e.g. from landslides or fire). Importantly, this review noted reduced viability of *K. kauaiensis* that was attributed to low census numbers.

Because our analyses were conducted on historically collected samples (circa 2000), our results reflect populations that may have been five-fold or greater in number than at present. Accordingly, the estimate of diversity presented here (π=1.29 x 10^-3^) is likely greater than the diversity that remains in these diminished populations. Importantly, however, the analyses here provide a perspective on diversity prior to the bottleneck experienced in the last two decades, allowing future sampling to uncover what proportion was lost. Our analyses suggest that *K. kauaiensis* consists of four genetic groups that largely correspond to their geographic areas. At the time of sampling, these populations generally exhibited more intrapopulation diversity than was found among all sampled individuals of either *K. cookei* or *K. drynarioides*, and genetic distance among individuals was far greater than found among individuals of either congener (Figure 2). Because these historical samples can be traced back to individual trees, the genetic framework presented here can be used to maximize genetic differentiation among outplants in the current managed populations.

## Conclusion

Island species are particularly at risk of extinction. On the Hawaiian archipelago, the *Kokia* genus has experienced dramatic reduction in habitat, mainly as a consequence of human activities, with one species already extinct and two out of three extant species nearly or completely gone from the wild. Currently, intensive conservation efforts are in place to save each species from extinction, leveraging managed populations in protected areas and captive propagation. Overall, our findings document low diversity and high homozygosity within each *Kokia* species, likely resulting in some amount of inbreeding depression whose outcomes are a concern for conservation (Kariyat & Stephenson, 2019). Importantly, our framework identifies hidden diversity in a previously misidentified sample, highlighting the value in surveying conservation materials. We believe that future conservation efforts will benefit from the results of this study, by informing the development of populations that may increase genetic diversity, and therefore robustness, in these populations.

## Acknowledgments

The authors thank Sierra McDaniel for providing materials and historical records regarding *K. drynarioides* and the HAVO *K. cookei*. The authors also thank Henry Oppenheimer for information regarding the history of *K. cookei*.

## Data availability

All code is available from https://github.com/Wendellab/KokiaResequencing. Sequencing data are deposited at NCBI in the SRA under PRJNA1229648.

## Benefit sharing

Benefits Generated: Benefits from this research include in-depth genetic information for individual trees and populations tracked by the Hawai‘i State Department of Forestry and Wildlife (DOFAW) and the Plant Extinction Prevention Program (PEPP). Although unreported here (due to the endangered nature), the exact location of each tree sampled is known to DOFAW and PEPP. The information provided here will be used by DOFAW and PEPP to inform conservation planning, diversify managed populations, and identify important sources of biodiversity for long-term preservation (i.e., seed banks).

## Author contributions

WN and EK conducted analyses, wrote the first draft, and generated figures; JFW contributed to the manuscript; CH, ZVM, and OP generated DNA libraries and supervised sequencing; MY and CWM provided DNA and provenance information; MJK and JVDM provided material and supervised sample collection; DGP acquired funding and provided supervision; MAA and CEG contributed to analyses, figures, and writing, and supervised the project. All authors reviewed and accepted the final version of the manuscript.

## Supplemental Figure Legends

*Supplementary Figure 1*

Interspecific and intraspecifc LEA analysis for three extant *Kokia* species, with sample information labeled.

*Supplementary Figure 2*

Species-specific PCA (A) and LEA (B) analysis for *K. cookei* including the newly identified HAVO tree (Kd_HV_1Ar). The HAVO sample is identified in the PCA using blue (versus red) coloring, reflecting its original species assignment.

*Supplementary Figure 3*

LEA populations by site for *K. drynarioides*, both for the interspecific and intraspecific analyses.

