## Supplementary material for "Population Genomics Informs Conservation Strategies for Critically Endangered *Kokia* Species in Hawai‘i": KokiaSupp_FigS1.pdf

Supplementary Figure 1

Interspecific and intraspecifc LEA analysis for three extant Kokia species, with sample information labeled.

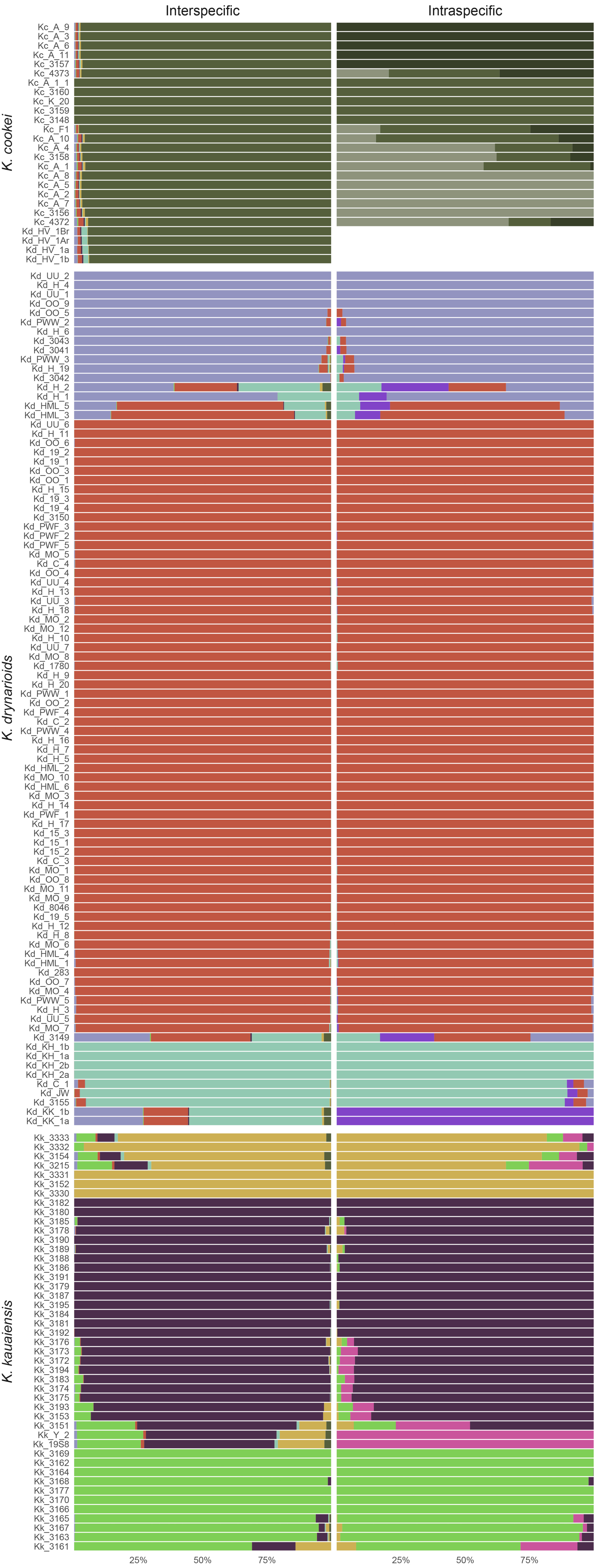
