## Supplementary material for "Population Genomics Informs Conservation Strategies for Critically Endangered *Kokia* Species in Hawai‘i": KokiaSupp_FigS2.pdf

Supplementary Figure 2
Species-specific PCA (A) and LEA (B) analysis for K. cookei including the newly identified HAVO tree (Kd_HV_1Ar). The HAVO sample is identified in the PCA using blue (versus red) coloring, reflecting its original species assignment.

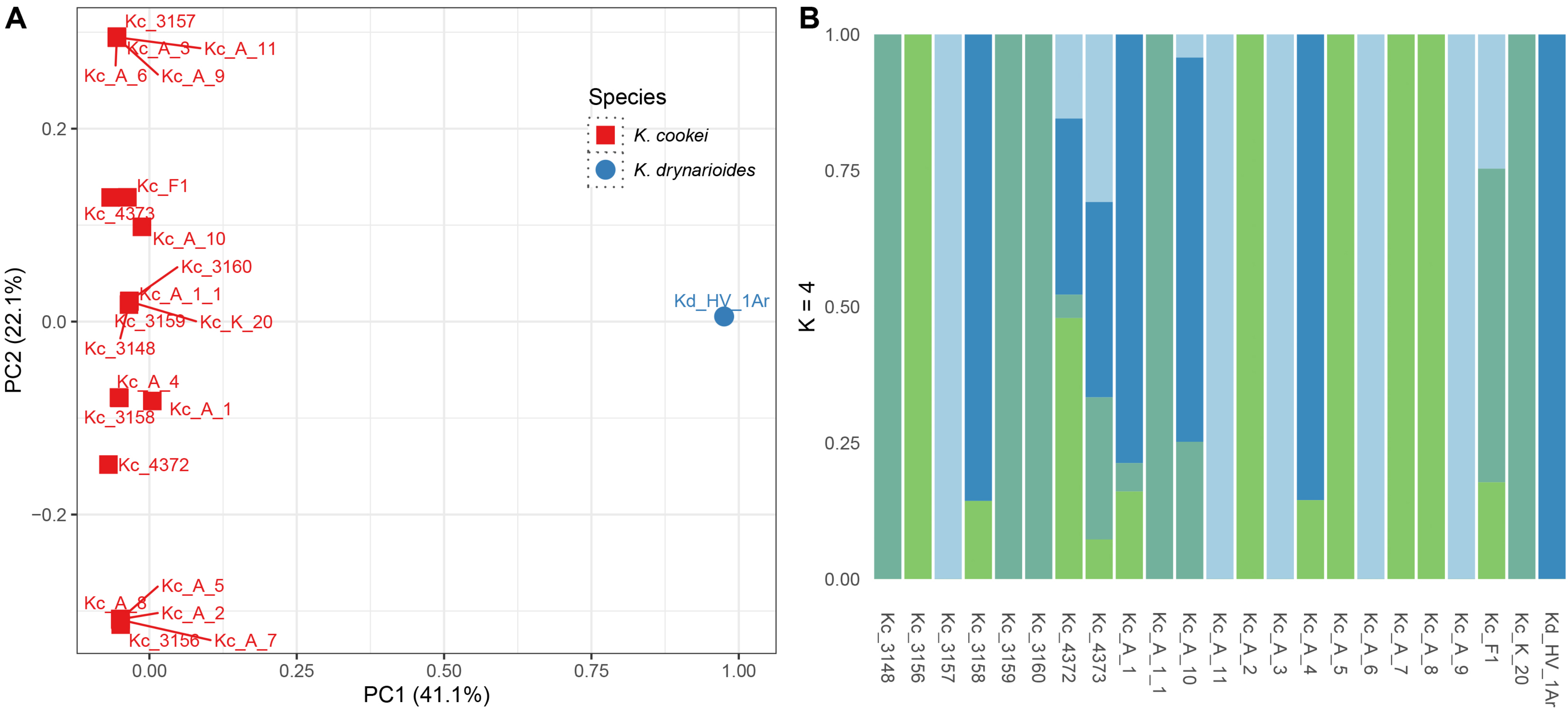
