## Supplementary material for "Population Genomics Informs Conservation Strategies for Critically Endangered *Kokia* Species in Hawai‘i": KokiaSupp_FigS3.pdf

Supplementary Figure 3

LEA populations by site for K. drynarioides, both for the interspecific and intraspecific analyses.

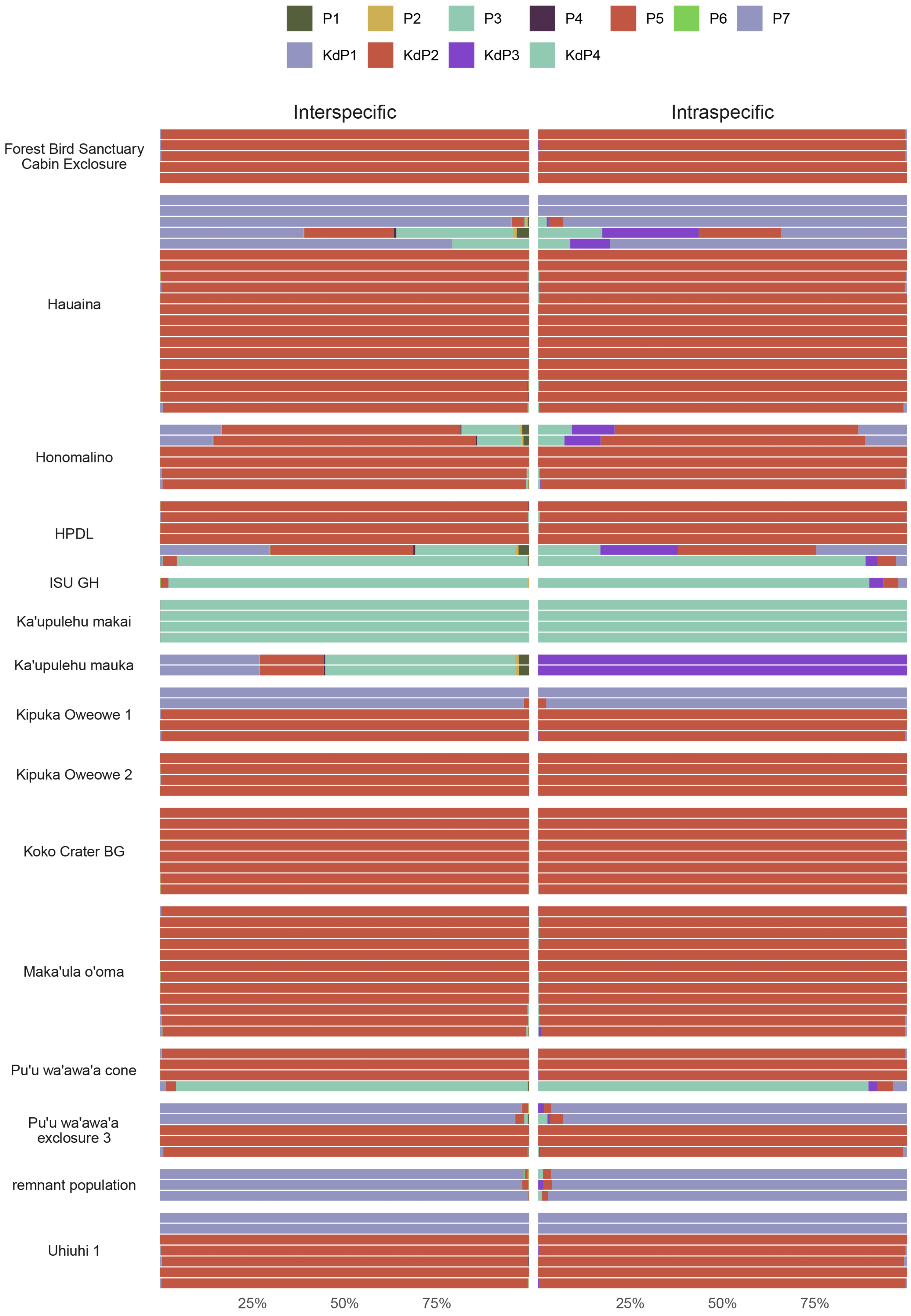
